# Cell Cycle Regulation of Merkel Cell Polyomavirus Replication and Genome Inheritance in Single Cells

**DOI:** 10.64898/2026.07.31.741607

**Authors:** Steven W. Smeal, Hongzhao Zhou, Md Musaddaqul Hasib, Bizunesh Abere, Gabriel J. Kowalczyk, David L. Schipper, Yufei Huang, Patrick Moore, Yuan Chang, Robin E.C. Lee

## Abstract

Merkel cell polyomavirus (MCV) establishes a near-ubiquitous, asymptomatic infection in humans but on rare occasions drives Merkel cell carcinoma, an aggressive skin cancer. How MCV replication is coordinated with host cell-cycle progression, and how this coordination shapes infected-cell fate, has remained unresolved. Here we combine reporter virus live-cell imaging, pharmacologic cell-cycle perturbation, flow cytometry, and single-cell transcriptomics to examine MCV replication dynamics. VP1 late gene expression occurs almost exclusively during late S or G2, is associated with prolonged G2 arrest, and frequently culminates in cell death. Strikingly, cells that divide before virus replication give rise to daughter cells that synchronously initiate viral replication and VP1 expression, revealing mitotic inheritance of silent viral genomes, a pattern seen for viruses with a known latency lifecycle. Single-cell RNA sequencing further identifies a transient antiviral and inflammatory response that is subsequently suppressed in surviving, cell cycle arrested cells. These findings define the cell-cycle logic, inheritance, and host consequences of MCV replication at single-cell resolution.

## Introduction

Merkel cell polyomavirus (MCV), discovered in 2008^1^, causes approximately 80% of Merkel cell carcinoma (MCC) cases, a rare but highly aggressive neuroendocrine skin cancer with a five-year disease-associated mortality approaching 50%^2^. The incidence of MCC has risen markedly in recent years, reaching over 3000 cases annually in the US, driven by aging populations and improved diagnosis, underscoring the clinical urgency of understanding MCV biology^3,4^.

Despite its oncogenic potential, MCV is a near-ubiquitous and asymptomatic human skin infection^5,6^. Tumorigenesis arises from a rare replication accident in which the viral genome acquires replication-disabling mutations and becomes clonally integrated into the host genome, resulting in constitutive expression of viral early oncoproteins—a dead end for the virus but a transformative event for the host cell^7–9^. Despite accumulating molecular insights, the cell-cycle stage at which MCV replication initiates and how replication alters infected cell fate remain unclear, particularly at single-cell resolution.

Like other polyomaviruses, the MCV genome is organized into early (ER) and late (LR) transcriptional regions separated by a non-coding control region (NCCR) that contains transcriptional regulatory elements and the viral origin of replication^1,10,11^. The ER encodes several tumor (T) antigen proteins generated through alternative splicing and recoding^1,9,12–14^ including large T (LT), small T (sT), and additional isoforms such as 57kT and middle T/alternative LT open reading frame (MT/ALTO). LT functions as a viral replicative helicase, initiating DNA replication by melting the replication origin and recruiting host replication machinery^15^. sT and other isoforms modulate LT stability, viral RNA processing, and host signaling pathways that influence viral replication and cell physiology ^16–19^. Initiation of viral DNA replication is required for late gene transcription, which encodes the VP1 and VP2 capsid proteins responsible for virion assembly and genome encapsidation^20^.

As a small DNA viruses, MCV relies on host cell cycle machinery to support replication^18,19,21,22^. Polyomaviruses typically manipulate the host cell-cycle regulators to create an environment favorable for viral DNA synthesis. However, the precise relationship between MCV replication and host cell-cycle progression remains incompletely understood. For example, related human polyomaviruses use distinct strategies to access replication-permissive states: BK virus induces prolonged S-phase through activation of the DNA damage response, whereas JC virus promotes S-phase entry through alternative regulatory mechanisms. Whether MCV uses similar or distinct strategies to coordinate viral replication with host cell cycle transitions remains unclear^23–25^. In addition, little is known about the fate of viral genomes in infected cells prior to the onset of replication. For some DNA viruses, viral genomes can persist in a transcriptionally silent state and be transmitted through mitosis before reactivation. Whether polyomavirus genomes exhibit similar behavior during early infection has not been explored.

Progress in addressing these questions has been limited by technical challenges. Like archetypal BKV and JCV polyomaviruses ^26,27^, wild-type MCV infects cells inefficiently in laboratory systems ^28,29^, limiting bulk studies of replication dynamics. Moreover, productive replication events are rare and asynchronous, making it difficult to connect viral gene expression with host cell responses using population-level approaches.

Here, we overcome these limitations using live-cell imaging to resolve MCV replication dynamics at single-cell resolution. We leverage a fluorescent MCV reporter virus expressing a VP1–mScarlet fusion protein together with fluorescent cell-cycle reporters, enabling direct visualization of the early lifecycle of this virus and host cell-cycle progression in living cells. Because VP1 expression is tightly coupled with MCV replication, we use VP1 expression as a surrogate marker for viral DNA replication^30,31^. In parallel, single-cell RNA sequencing (scRNA-seq) allows us to define host transcriptional responses associated with early and late viral gene expression. Using these complementary approaches, we establish how MCV replication couples to host cell-cycle progression. We find that viral replication initiates within a restricted S/G2 window, induces prolonged G2 arrest, and frequently culminates in cell death. Strikingly, when infected cells progress through mitosis before replication begins, viral genomes are transmitted to daughter cells that subsequently initiate replication synchronously, revealing mitotic inheritance of replication-competent viral genomes. Consistent with prior studies of polyomavirus infection, single-cell transcriptomic analysis also detects a transient antiviral transcriptional response during replication that is subsequently attenuated in surviving, cell-cycle arrested cells. By quantifying viral protein expression and host dynamics directly, our work provides new understanding of how tumorigenic DNA viruses persist, propagate, and drive infected cell fates and disease.

## Results

### Single-Cell Imaging Platform Resolves MCV Replication Timing and Cell Cycle State

To relate the timing of MCV replication to cell-cycle stage in single U2OS cells, we classified cells as G1 or S/G2/M using a Geminin(1/110) far-red fusion reporter that accumulates during S/G2/M and is degraded in G1^32^. Single-cell DNA content was measured by adding trace concentrations of Hoechst 33342 immediately before imaging (see^33,34^ and Methods). Integrated nuclear Hoechst intensity increases during S phase and plateaus in G2, allowing real-time discrimination of cell cycle states ^33,35^. Together, these reporters provided reliable cell detection and classification of G1, S, and G2 phases (Fig. 1A-C).

**Fig. 1:**
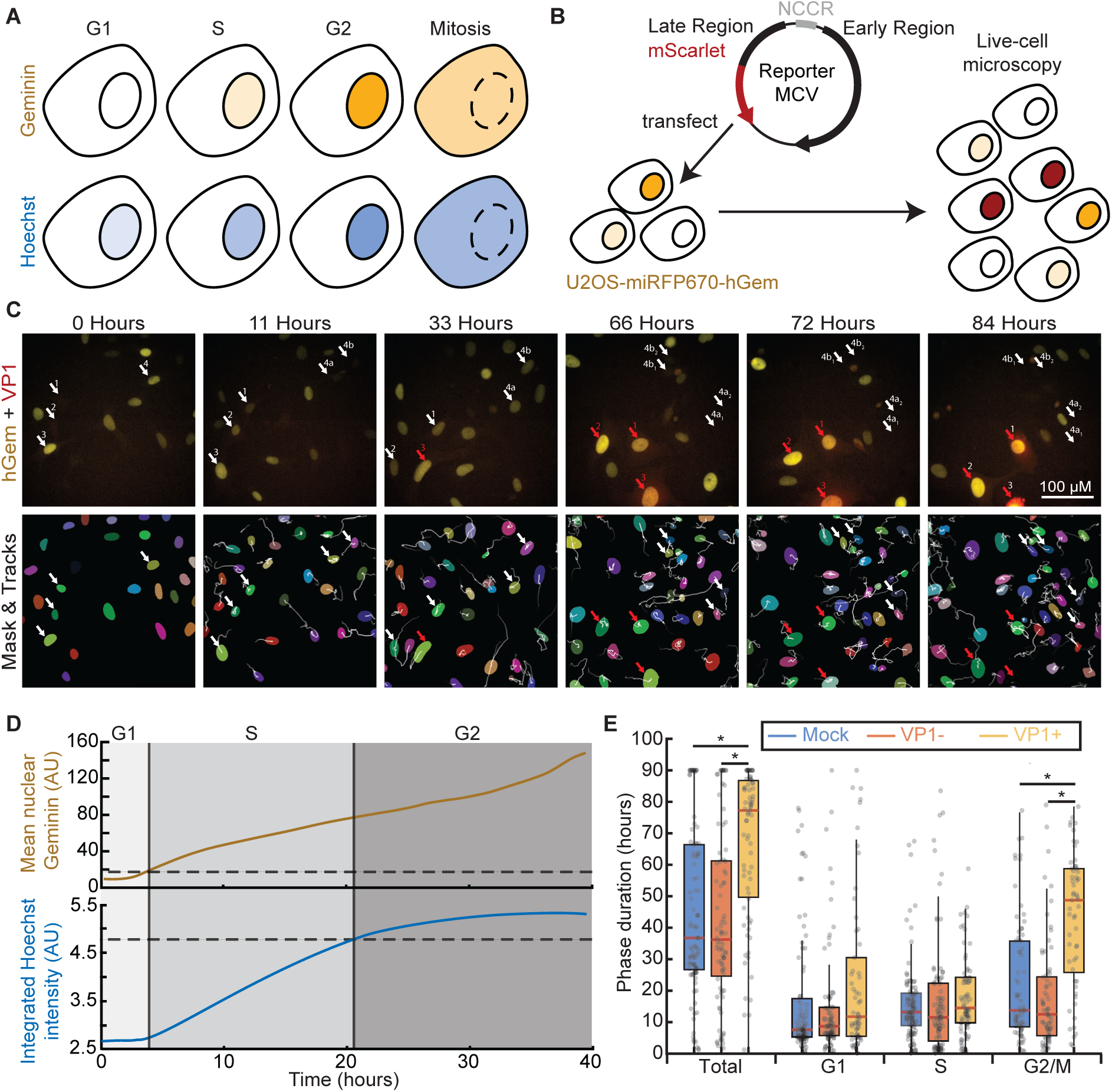
U2OS cells transfected with an MCV reporter and tracked by live-cell imaging. (A) Schematic of cell cycle phase determination using two measurements: mean nuclear fluorescence intensity of the Geminin reporter (miRFP670-hGem(1–110)) to indicate entry into S phase (top), and nuclear Hoechst staining to measure total DNA content, which increases during S phase and plateaus at G2 (bottom), enabling accurate identification of cell cycle stage during imaging. (B) U2OS cells transfected with MCVmc construct encoding VP1 fused to mScarlet, enabling visualization of VP1 accumulation during live-cell imaging. (C) Top: Time-lapse images of U2OS cells transfected with MCVmc and stably expressing the Geminin reporter. The Geminin signal (yellow) was merged with MCV reporter (red) to track cell cycle phase and VP1 expression over time. Bottom: Segmentation masks and cell tracks, shown as gray trailing lines, corresponding to the live-cell imaging in the top panel. White arrows show cells that are VP1(-), red arrows are VP1(+). Number labels represent the same cell across images. 4a and 4b are daughter cells of cell 4, 4a_1_ and 4a_2_ are daughter cells of 4a, and 4b_1_ and 4b_2_ are daughter cells of 4b. (D) Schematic illustrating how mean nuclear Geminin and integrated Hoechst intensity were used to assign cell cycle phase. Entry into S phase (top) was defined by Geminin levels rising above background (dashed line), where background was determined as the 99th percentile of Geminin-negative cells. Entry into G2 phase (bottom) was defined when integrated nuclear DNA content reached 90% of the maximum value (dashed line). (E) Boxplot showing the duration of each cell cycle phase in mock, VP1(–), and VP1(+) cells. Asterisks indicate statistical significance (Student’s t-test,* p < 10^-2^).

To monitor viral replication, U2OS reporter cells were transfected with MCVmc minicircle genome DNA, incubated for 72 hours (before which VP1 expression is not detected by western blot^36^), and then imaged over multiple days (Fig. 1B). Fluorescence images were acquired every 15 minutes across Hoechst, Geminin, and VP1 channels (Fig. 1C, top). Nuclei were segmented from the Hoechst channel and tracked using machine-learning algorithms with manual curation (Fig. 1C, bottom; see Methods), yielding multi-channel single-cell trajectories.

Cell-cycle phases were assigned from single-cell trajectories using a combination of fluorescence signal thresholds for nuclear Geminin and Hoechst (Fig. 1D). Cells lacking Geminin expression, defined as having mean nuclear fluorescence intensities below the 99th percentile of Geminin-negative cells (Supplementary Fig. 1A), were classified as G1, whereas Geminin-positive cells were separated into S or G2 phases based on DNA content dynamics. During S phase, the integrated fluorescence of nuclear Hoechst reflects DNA replication and increases approximately twofold before reaching a plateau in G2. Based on this pattern, the S-to-G2 transition was defined when nuclear DNA content reached 90% of its maximum value (Fig. 1D, bottom). This approach enabled reconstruction of single-cell cell-cycle trajectories and phase durations.

### MCV VP1 Late Gene Expression Initiates in S/G2 and Prolongs G2/M

Cell-cycle dynamics were compared between mock-transfected cells and MCVmc-transfected cells. Transfected cells were classified as VP1-expressing (VP1+) or VP1-negative (VP1-) based on single-cell trajectories exceeding the 99th percentile VP1 signal observed in mock cells (Supplementary Fig. 1B). VP1+ cells exhibited a significantly longer total cycle duration than mock or VP1-cells (Fig. 1E; p < 10^-2^, Student’s t-test). Although some VP1+ cells showed prolonged G1 phases, these differences were not statistically significant. Instead, the increase in cell-cycle duration was primarily driven by a prolonged G2 phase (p < 10^-2^, Student’s t-test), indicating that VP1 expression extends G2 or delays mitotic progression.

Consistent with bulk measurements of MCV replication kinetics, these single-cell results indicate that MCV DNA replication and late gene expression occur predominantly within cells in a G2-like state. Cell death was also elevated among VP1+ cells (27%) compared with mock (9%) (p = 0.002, Fisher’s exact test) and VP1-cells (14%) (p = 0.048, Fisher’s exact test) (Supplementary Fig. 1c), linking productive replication to both cell-cycle arrest and increased cell mortality.

### Replication Timing/VP1 Expression and Infected Cell Fate: Prolonged Arrest or Cell Death

Single-cell kymographs were used to visualize trajectories of Geminin, Hoechst and VP1 fluorescence for mock and VP1-cells (Supplementary Fig. 2A, B) or VP1+ cells (Fig. 2B). Each trajectory was classified into one of four fate classes based on observed mitotic birth and terminal fate (mitosis or death; Fig. 2A). Trajectories within each class were ordered by entry into S phase (right-pointing arrow), with G2-phase (left-pointing arrow), mitosis (green circle), or cell death (red X) as indicated.

**Fig. 2:**
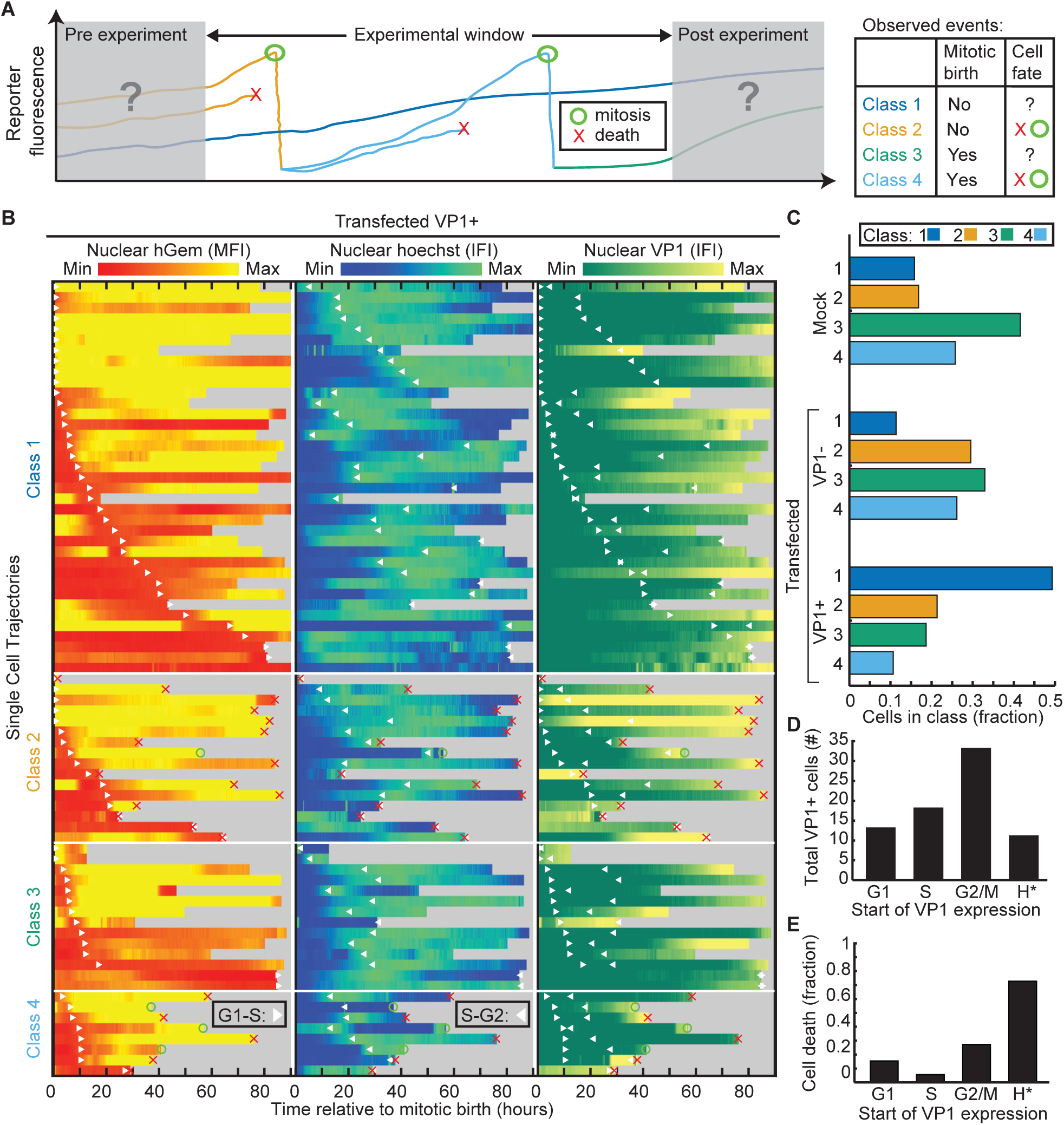
VP1(+) cells show a higher proportion of Class 1 trajectories and initiate VP1 accumulation during S or G2 phase. (A) Single cell trajectories were classified into four groups based on observed events during imaging. Classes 1 and 2 lacked an observed mitotic birth event, whereas Classes 3 and 4 included one. Classes 1 and 3 ended with unknown fates (experiment ended or the cell left the frame), while Classes 2 and 4 had an observed fate, either death (X) or mitosis (O). (B) Kymographs of VP1(+) U2OS cell trajectories, separated into the four classes. Within each class, trajectories are aligned by the start of S phase. Each trajectory displays mean fluorescence intensity (MFI) of nuclear Geminin, integrated fluorescence intensity (IFI) of nuclear Hoechst, and nuclear VP1 IFI over time. Rightward arrows mark the start of S phase, and leftward arrows mark the start of G2, shown as white in the Geminin MFI or Hoechst IFI plots, and as red or blue, respectively, in the VP1 IFI. (C) Bar graph showing the distribution of cell classes within each group (Mock, VP1(–), and VP1(+)), with fractions normalized to sum to 1. (D) Bar graph showing the cell cycle stage at which VP1 begins to accumulate in VP1(+) cells. H* indicates trajectories with high VP1 levels present at the start of the cell track. (E) Bar graph showing the fraction of cells that died, grouped by the cell cycle stage at which VP1 accumulation began. H* indicates trajectories with high VP1 already present at the start of the cell track.

Class 1 cells were not observed to undergo mitosis or cell death. These cells tended to be cell cycle arrested cells since the average cell cycle duration for uninfected is 35 hours for U2OS cells. This class, however, also included cells with shortened trajectories due to cells entering or exiting the field of view. Class 2 cell contained those having a known fate (mitosis or cell death). Class 3 consisted of cells having an observed mitotic birth but no defined fate. Finally, Class 4 included cells in which both mitosis and fate were determined. This approach assigned each trajectory to a single class and enabled direct comparisons of VP1 accumulation and cell-fate dynamics while accounting for limitations imposed by experimental parameters, such as imaging duration and field of view.

Mock and VP1-cells showed similar class distributions (Fig. 2C) with Class 1 (unknown birth, unknown fate) representing the smallest fraction. By contrast, VP1+ cells predominantly fell into Class 1 (Fig. 2C) and these cells show a prolonged S/G2 Geminin positivity (Supplementary Fig. 2C; p < 0.05, Student’s t-test). Cells with observed mitotic birth (classes 3 and 4) were also depleted in VP1+ cells.

VP1 expression onset was defined by visual inspection as the earliest time point of sustained accumulation above the same cell’s baseline. Our analysis revealed that VP1 accumulation initiates primarily during S and G2 phases (Fig. 2D). A subset of cells exhibited high VP1 content at the start of the imaging experiment (H* cells), with ∼70% undergoing cell death versus ∼30% of cells that initiated VP1 accumulation in G2/M (Fig. 2E). This strong association indicates that rapid and sustained VP1 expression following virus replication predisposes cells to death, potentially through viral protein overload or replication-induced stress responses. Class 1 and 3 VP1+ trajectories are therefore likely to represent a continuum progressing toward death as observed in Class 2 cells, rather than remaining in an undefined state. This implies that MCV replication not only inhibits mitotic entry but also prolongs cell-fate decisions, as many VP1+ cells persist in an extended G2-like state for multiple days before ultimately committing to cell death.

### Cells that Divide Prior to Replication Transmit Silent MCV Genomes to Daughter Cells

A subset of 12 cells were initially VP1-, underwent mitosis, and then began expressing VP1. Capturing this transition required identifying cells that divided before the onset of detectable VP1 expression and then tracking both daughters through the multi-day window over which replication initiates—a conjunction of two comparatively rare events that yielded 12 informative parent-cell lineages across our imaging cohort. Despite this modest number, the paired design, in which each pair of daughters shares an identical genetic and immediate-environmental history, provides a high-resolution, within lineage comparison that is not accessible by population-level or fixed-cell approaches. This cohort of dividing VP1-parent cells with VP1+ daughters (subset of classes 3 and 4) enabled sister-cell measurements to compare the timing of VP1 accumulation (Fig. 3a). When both sister cells were observed to express VP1, we defined the time of VP1 onset relative to mitotic birth for the early-expressing sister (t_d1_) and plotted it against the time of VP1 onset for the later-expressing sister (t_d2_; Fig. 3B, blue data points). When both sisters expressed VP1, symmetric behavior was observed in 7 of 8 sister pairs, with VP1 onset occurring within 10 hours of each other. Given the small number of informative pairs (n=8 with detectable VP1 in both daughters), we report this concordance descriptively (Fig. 3B and C), rather than as a population-level estimate of penetrance or timing variance; the consistency of the symmetric pattern across nearly all pairs nonetheless argues against this being a chance occurrence. There was limited occurrence of asymmetric sister-cell behavior, where VP1 expression was either delayed by greater than 10 hours or not observed in the sister (Figs. 3B and C, and Supplementary Fig. 3).

**Fig. 3:**
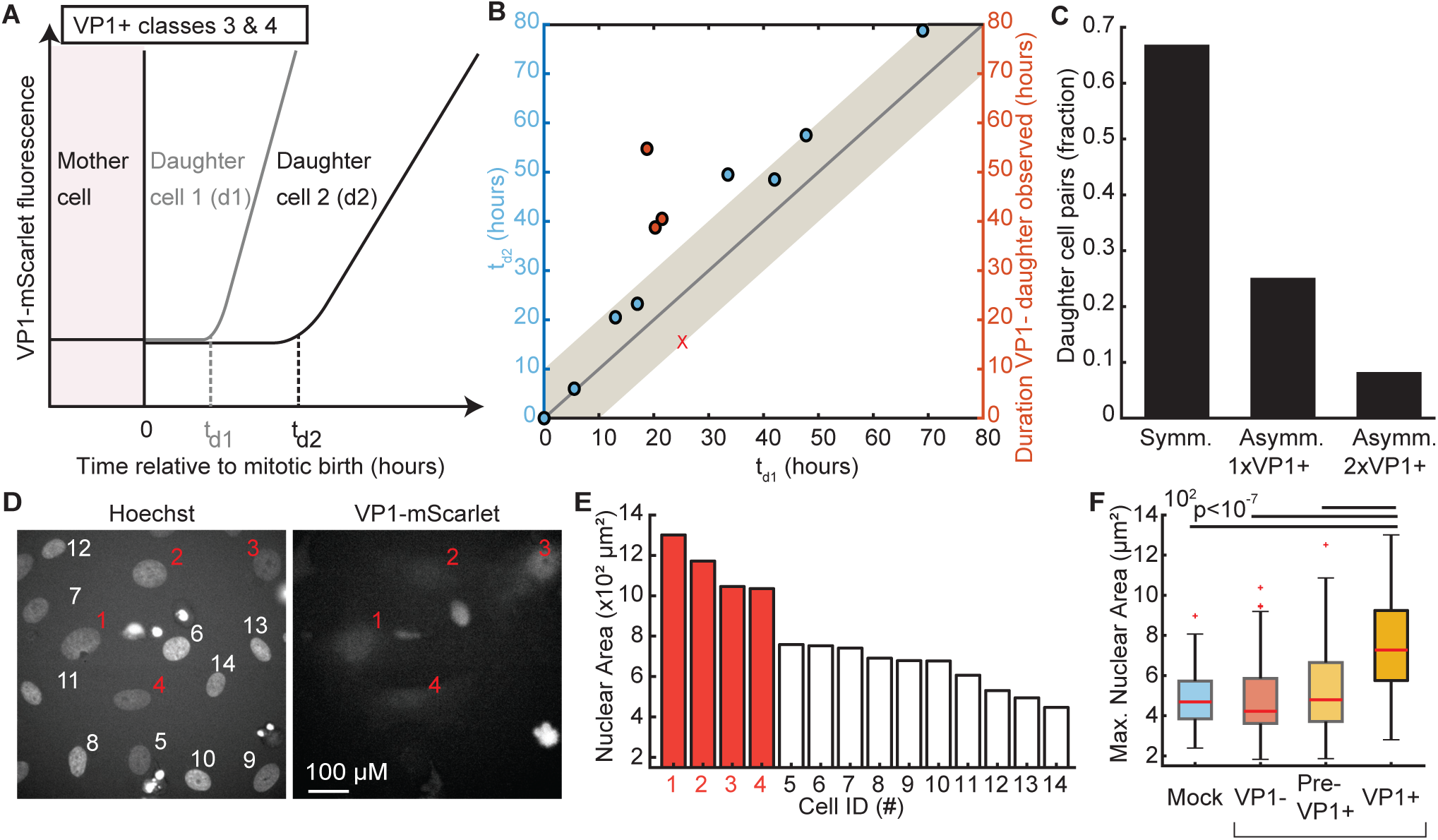
Sister cells show predominantly symmetric VP1 accumulation within a 10-hour window. (A) Schematic illustrating VP1 accumulation timing in daughter cells. Accumulation onset is measured as the time from mitotic birth to detectable VP1 expression in the first (t_d1_) and second (t_d2_) daughters. (B) Dual-axis plot with the left axis (blue) showing the time from mitotic birth at which daughter 1 and daughter 2 initiated VP1 accumulation. Daughter 1 was defined as the first cell to express VP1. The right axis (red) corresponds to cases where only one daughter expressed VP1, representing the total duration of the observed VP1 trajectory. Red X marks indicate daughter cells that left the imaging frame. The gray region highlights a ±10-hour window between the accumulation times of daughter 1 and daughter 2. (C) Bar graph showing the distribution of VP1 accumulation patterns between sister cells. Categories include symmetric expression (both daughters accumulate VP1 within 10 hours), asymmetric expression where only one daughter accumulates VP1 (Asymmetric 1x VP1(+)), and asymmetric timing where the second daughter accumulates VP1 outside the 10-hour window (Asymmetric 2x VP1(+)). (D) Example image of Hoechst-stained nuclei and corresponding VP1-mscarlet image illustrating variation in nuclear size and VP1 positivity. (E) Bar plot of nuclear areas corresponding to the nuclei shown in panel A, with red numbers denoting VP1(+) cells. (F) Boxplot of maximum nuclear area for Mock, VP1(–), Pre-VP1+ and VP1(+) cells. Pairwise differences were assessed using two-sample t-tests. Pre-VP1+ expression only considered the single cell trajectory until VP1 started expression.

There were 4 sister-cell pairs where only one cell expressed VP1, in which case we defined t_d2_ as the total duration for which the sister cell was observed before the trajectory ended, either due to lost tracking or the end of the experiment (Fig. 3B, red data points). The symmetric fates of most sister pairs suggest that these mother cells either maintained multiple viral genomes that were passaged to both daughters, or that viral DNA replication occurred prior to division, but VP1 expression remained undetectable at the time of mitosis due to the delay between viral DNA replication and protein accumulation. VP1-associated mitotic arrest or death only arose following mitosis, when detectable VP1 expression emerged in both daughter cells.

### Nuclear enlargement occurs after VP1 expression

We observed that VP1+ cells often appeared to have enlarged nuclei relative to VP1- and mock-transfected cells (Fig. 3D). Quantitative analysis confirmed that the nuclear area is significantly increased following VP1 expression (Fig. 3E). We next compared the nuclear area of VP1+ cells at time points that precede detection of VP1 fluorescence, termed “pre-expression” cells, to ask whether nuclear enlargement occurs before viral DNA replication. We did not observe a significant increase in nuclear area in pre-expression cells relative to uninfected cells; however, nuclear area increased significantly between pre-expression and post-expression (VP1+) cells (Fig. 3F; p<10^-7^). Together, these results indicate that MCVmc carrying cells can reside in a silent state, with the onset of VP1 expression coinciding with nuclear alterations and morphological changes.

### Cell-Cycle Perturbations Confirm Preferential MCV Replication in G2/M phase

To validate our live cell imaging results with orthogonal approaches, we examined the relationship between MCV replication and cell-cycle phase using pharmacologic cell-cycle perturbations combined with flow cytometry. U2OS cells were transfected with MCVmc or mock (reagent-only, no MCV DNA) and, three days later, treated for 24 hours with cell-cycle arrest agents or DMSO control: 0.5 mM mimosine (G1 arrest), 10 µM aphidicolin (S-phase arrest), or 9 µM RO-3306 (G2 arrest). After arrest, cells were then released into fresh medium and harvested at 0, 6, 12, and 24 hours post-release. Cell cycle profiles were determined by bromodeoxyuridine (BrdU) incorporation and FxCycle Violet DNA staining (Supplementary Fig. 4A).

In asynchronous DMSO-treated cultures, the majority cells were in G1 phase (59.2%), followed by S phase (27.9%) and G2 phase (12.8%) (Fig. 4A, Fig. S4A). Fewer than 5% of cells were VP1 positive under these conditions or following G1 arrest with mimosine. VP1 positivity cells increased modestly in aphidicolin-treated cells (∼8%) and was highest in RO-3306–treated cultures (∼24%; Fig. 4B), indicating enrichment of late viral gene expression when cells were arrested in G2. To quantify this relationship between infection and cell-cycle phase, we calculated the conditional probability of VP1 positivity for each cell-cycle phase under all treatment conditions. In every case, cells with late S/G2 DNA content showed the highest probability of VP1 expression (Supplementary Fig. 4B).

**Fig. 4:**
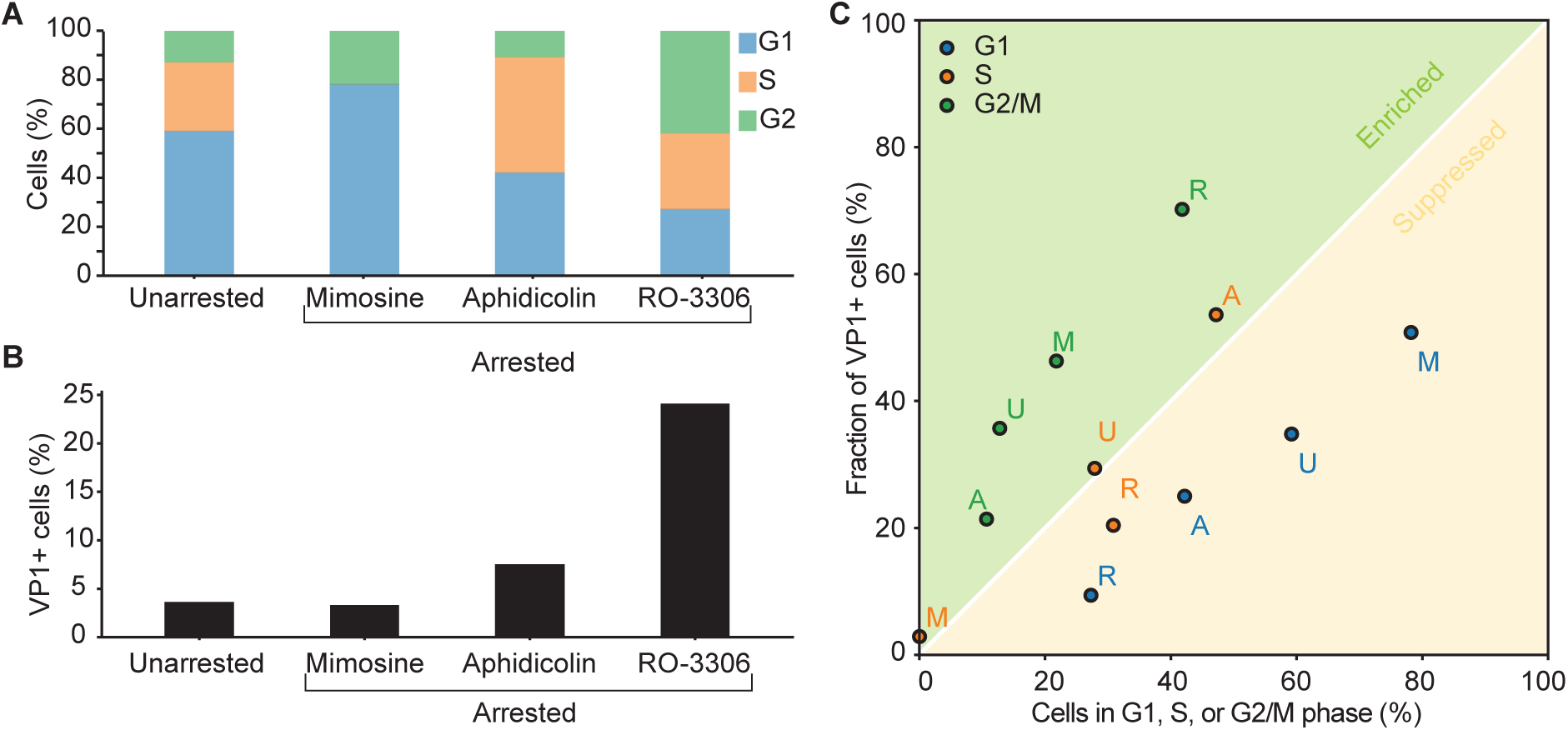
Flow cytometry reveals VP1-positive cells are enriched in the G2 phase. (A) Percentage of cells in G1, S, or G2 phase after 24 hours of treatment with cell cycle arrest agents: mimosine (G1 arrest), aphidicolin (S phase arrest), and RO-3306 (G2 arrest). (B) Percentage of VP1-positive cells following treatment with cell cycle arrest agents: mimosine (G1 arrest), aphidicolin (S phase arrest), and RO-3306 (G2 arrest). (C) Scatter plot comparing cell cycle distribution of the total population with that of VP1(+) cells by phase. The x-axis shows the fraction of cells in G1, S, or G2/M; the y-axis shows the fraction of VP1(+) cells within the corresponding phase. Points are color-coded by phase and labeled by treatment: U (unarrested), M (mimosine), A (aphidicolin), and R (RO-3306).

Under a neutral model, where VP1 expressions occur equally across cell cycle phases, the fraction of VP1 positive cells would scale directly with the total fraction of cells in each phase, landing on the perfect diagonal line in this space (Fig. 4C, white line). Comparison of VP1+ cells relative to DNA content for unarrested (U), mimosine (M), aphidocolin (A), and RO-3306 (R) treated cells showed deviations from the neutral model (Fig. 4C). Regardless of the arrest conditions, cells with G1 DNA content (blue dots) were depleted, S-phase (orange dots) were approximately neutral, and G2/M cells (green dots) enriched for VP1 expression. Following release from arrest, VP1 expression increased as synchronized populations progressed through S phase and into G2 (Supplementary Fig 4C). These results mirror the single-cell imaging analysis and indicate that late MCV gene expression is strongly biased toward late S and G2 phases.

### MCV Infection Recapitulates Cell-Cyle-Restricted Replication and Nuclear Remodeling

To confirm the findings from MCVmc transfection studies (Fig 2-4), MCVmc reporter minicircle DNA was packaged into wild-type VP1/VP2 capsids and used to infect BJ-hTERT fibroblasts stably expressing the miRFP670-hGem at MOI = 10⁴ (see ref ^30^) and live cell imaging was performed at day 5 and 7 post infection (PI). In parallel wild type BJ-hTERT cells infected with the non-reporter MCV.mc Wt virus or mock infected were collected at day 5 and 7 PI for single cell sequencing (Fig. 5A). In imaging experiments, cells were classified as either G1 or S/G2/M based on the Geminin(1/110) far-red fusion reporter. Due to incomplete single-cell trajectories, caused by long replication times, increased cell mobility and overlapping neighboring cells we could not further classify the cells into S and G2/M phases. scRNA sequencing enabled quantification of host transcriptional states together with MCV early region (ER) and late region (LR) transcripts to assign cells into VP1 status as well as G1, S or G2/M phases of the cell cycle.

**Fig. 5:**
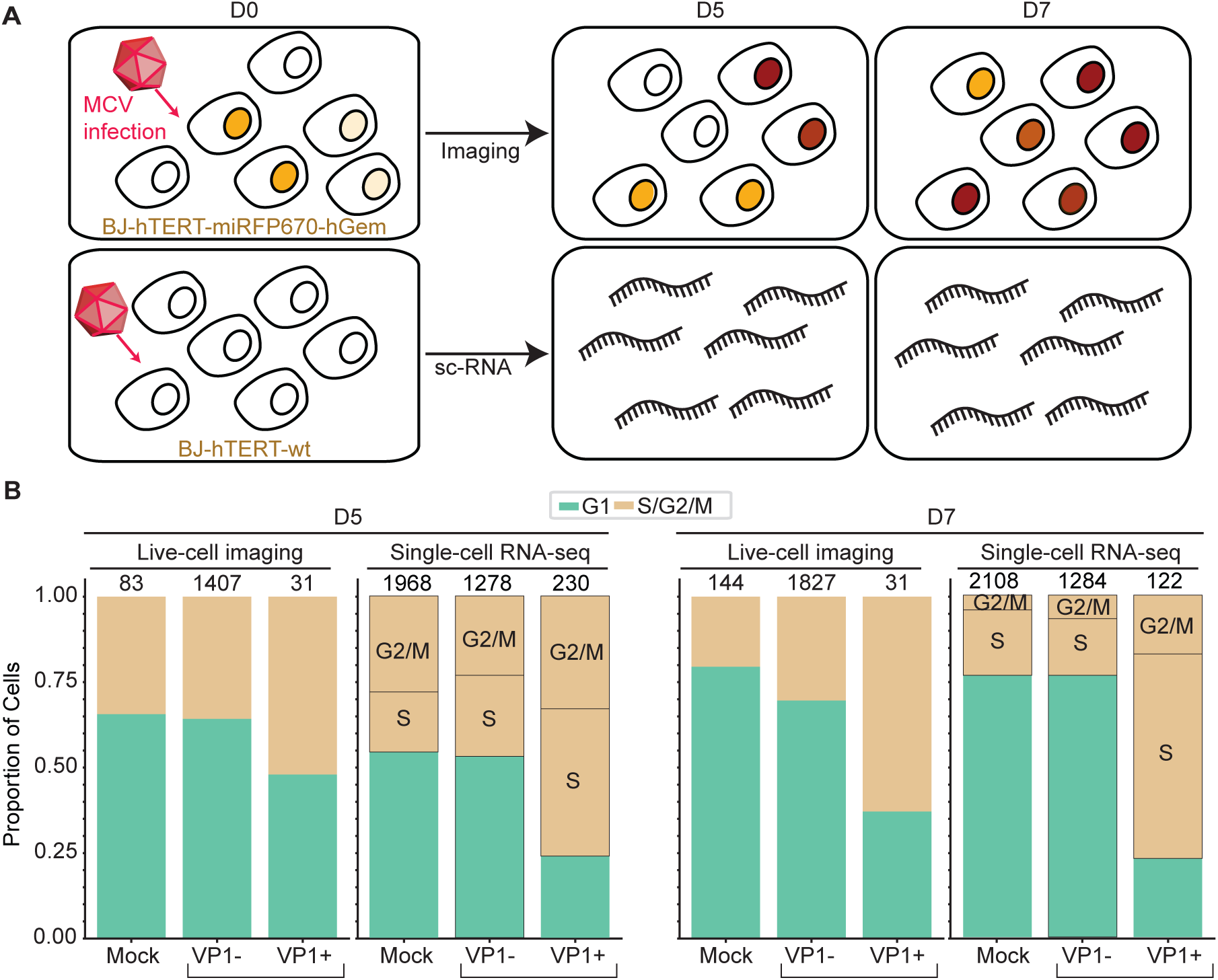
BJ-hTERT cells infected with MCV reveal an increased proportion of cells in S/G2/M in VP1(+) cells in both live cell and single cell RNA sequencing experiments. (A) Schematic of the experimental workflow. Bj-hTERT cells stably expressing the miRFP-670-hGEM reporter (for live cell imaging) or wt cells (for scSequencing) were infected with MCVmc reporter or MCVmc.Wt virus or mock infected. At day 5 or 7 post-infection (PI) for imaging or at Days 5 and 7 for scSequenceing, cells were either imaged or collected for single cell RNA sequencing. (B) Fraction of cells in G1 or S/G2/M phase at days 5 and day 7. For the live imaging, cell cycle phase was assigned based on the mean nuclear intensity of the Geminin reporter. For single cell RNA sequencing, cell cycle assignment was further refined into S, and G2/M phases based on the single cell sequencing results. VP1(-) cells were classified as being the combined bystander and cells expressing early response genes so the figure was directly comparable to the live cell imaging data.

Consistent with U2OS transfection experiments, VP1+ BJ-hTERT cells displayed larger nuclei (Supplementary Fig. 5A) and an enrichment in S/G2/M at days 5 and day 7 compared to mock-infected or VP1-cells (Fig. 5B). scRNA sequencing confirmed enrichment of S- and G2/M-phase gene signatures in infected cells (Fig. 5B and Supplementary Fig. 5B), consistent with cell-cycle arrest and senescence-associated nuclear remodeling^37^. Together, these orthogonal approaches demonstrate that VP1+ BJ-hTERT cells are enriched in S/G2 phases, reinforcing that VP1 expression induces S/G2 arrest.

### Single-Cell Transcriptomic Analysis Reveals Distinct Host Transcriptional Responses to MCV Infection

Single-cell transcriptomic analysis of BJ-hTERT cells at Days 3, 5, and 7 post-infection further defined the temporal dynamics of infection. UMAP analysis revealed at Day 3, mock and cells from corresponding infected sample were largely overlapping (cluster 1), indicating minimal early transcriptional divergence (Fig. 6A). By days 5 (cluster 2) and 7 (cluster 3), MCV-infected cells segregated into a pair of related clusters distinct from mock populations (clusters 2 and 3 versus cluster 4, Fig.6A). Based on viral transcript detection, cells were classified as mock (uninfected samples, no viral expression), bystander (infected culture, no detectable viral transcripts), ER-only (early region transcripts, not late genes), and ER+LR (both early and late region transcripts). At Day 5 PI, viral infection peaked with 24.7% (9.4% ER-only and 15.3% ER+LR, Fig. 6B, 6C) which declined to 13.5% (4.8% ER-only and 8.7% ER+LR) by D7 PI, likely reflecting loss of infected cells, consistent with elevated cell death observed by live-cell imaging.

**Fig. 6:**
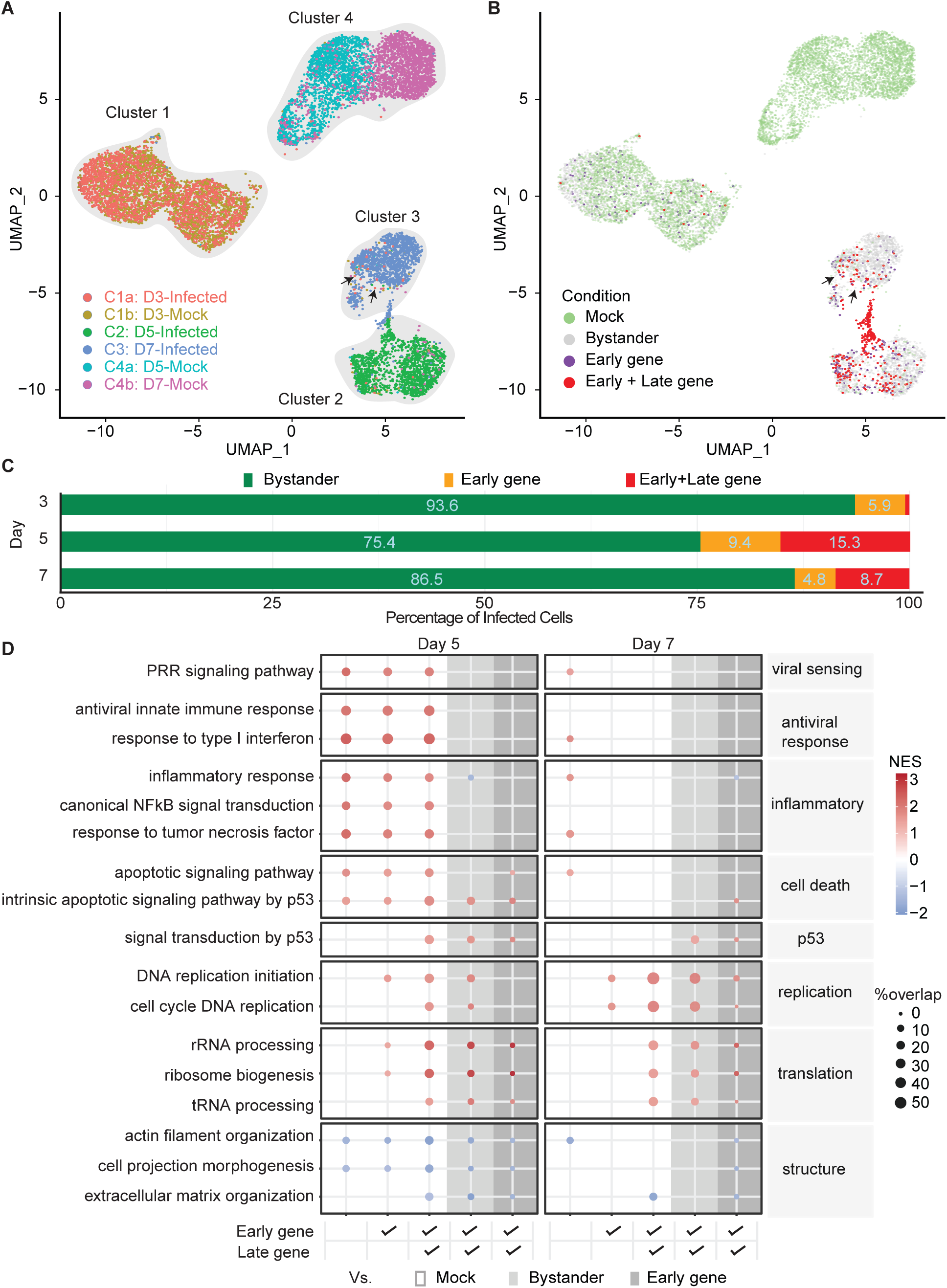
Single cell Bj-hTERT analysis shows distinct transciption profiles and reduced cell-shape regulation in response to MCV infection. (A) UMAP visualization of single cell RNA sequencing data from six conditions: mock and infected cells at 3, 5, and 7 days. Axes show UMAP_1 and UMAP_2. (B) UMAP visualization as in (A), with cells color-coded as mock, bystander (infected without viral mRNA expression), early gene (early viral gene expression only), and late gene (late viral gene expression). (C) Percentage of infected cells that are bystander, early, or late gene expressing cells. (D) GO analysis of the enriched functions (Padj < 0.05) in cells infected with MCV compared to mock. Color indicates normalized enrichment score; % overlap denotes the proportion of differentially expressed genes (log2FC ≥ 1, Padj < 0.05) associated with each function.

Differential gene expression analysis identified broad transcriptional responses across different cell states. ER+LR cells showed the largest divergence from mock at both Day 5 and Day 7 (Day 5 PI: 1288 up, 1143 down; Day 7 PI: 1809 up, 473 down, adjusted p < 0.05, |log2FC| ≥ 1, Supplementary Fig. 6A) likely due to active viral DNA replication induced stress. Bystander cells also showed substantial transcriptional changes relative to mock despite lacking detectable viral gene expression (Day 5 PI: 840 up, 730 down; Day 7 PI: 501 up, 476 down), potentially reflecting paracrine signaling from infected neighbors or the presence of transcriptionally silent viral genomes.

To identify transcriptional programs specifically attributable to complete viral replication beyond what the infected-culture environment or early viral gene expression alone explains, we directly compared ER+LR cells to bystander and ER-only populations. Direct comparisons revealed that ER+LR cells acquired an expanding set of transcriptional programs relative to bystander cells from Day 5 PI(335 up, 225 down) to Day 7 PI (956 up, 130 down), while differing from ER-only cells by far fewer genes at both timepoints (Day 5 PI: 60 up, 62 down; Day 7 PI: 67 up, 49 down)(Supplementary Fig. 6A). Overall, number of downregulated genes are decreased at day 7 PI relative to day 5 PI across all comparisons. Pathway enrichment analysis across these comparisons identified two broad categories of host transcriptional response distinguished by whether they require viral gene expression (Fig. 6D): a tissue-wide response shared across all infected-culture cells, and a set of responses specifically requiring viral gene expressions.

The first category comprised a tissue-wide antiviral and inflammatory response shared across bystander, ER-only, and ER+LR cells relative to mock significantly enriched at Day 5 PI across all infected-culture populations regardless of viral gene expression state (Fig. 6D). By Day 7 PI, these pathways were substantially diminished or absent across all populations, indicating near-complete resolution of the acute antiviral and inflammatory response. Cell death pathways showed a similar tissue-wide pattern at Day 5 PI with additional enrichment specifically in ER+LR cells relative to both bystander and ER-only populations, suggesting complete viral replication selectively amplifies the apoptotic arm of the stress response. Conversely, structural organization pathways were significantly downregulated in ER+LR cells at both timepoints, aligning with the nuclear enlargement and morphological changes observed by live-cell imaging, (Fig. 3F and supplementary Fig. 5A). The second category comprised transcriptional programs enriched in ER+LR or ER only but absent in bystander cells relative to mock, identifying programs that specifically require viral gene expression for activation. Replication pathways were enriched at both time points and amplified substantially by Day 7 PI, indicating sustained engagement of host replication machinery. A comprehensive translational reprogramming program was strongly and specifically elevated in ER+LR cells at both days, consistent with the protein synthesis demands of viral capsid assembly. Detailed gene-level analysis of each comparison and pathways are provided in (**Supplementary Table 1**).

To identify the upstream transcriptional regulators driving these pathway changes, we applied unbiased transcription factor (TF) activity inference to all cell states and timepoints. Direct comparison of TF activity between Day 5 PI and Day 7 PI within ER+LR cells confirmed that IRF9, STAT2, and IRF1 are the upstream drivers of these antiviral pathways showed the largest activity decline across the entire dataset (fold changes of 4-5) (supplementary Fig. 6B), providing mechanistic explanation for antiviral pathway resolution. Conversely, E2F2 and E2F3, which drive DNA replication initiation and cell cycle replication pathways enriched when compared to mock cells (supplementary Fig. 6C), but showed no significant change between Days 5 and 7 within ER+LR, indicating the replication program is constitutively maintained and that the progressive amplification of replication pathway enrichment from Day 5 to Day 7 PI reflects accumulating viral DNA replication activity driven by constitutively active E2F factors.

Together, these findings define the transcriptional state of MCV-replicating ER+LR cells as progressively diverging from both the infected-culture environment and early gene expression alone. At Day 5, ER+LR cells are dominated by antiviral signaling that resolves by Day 7 as IRF9, STAT2, and IRF1 activity declines, while viral gene expression-specific programs - while the replication transcriptional program, sustained by constitutively active E2F factors, progressively deepens. The selective amplification of apoptotic signaling specifically in ER+LR cells provides a transcriptional basis for the cell fate spectrum documented by live-cell imaging, collectively defining ER+LR cells as a progressively reprogrammed viral factory operating under sustained checkpoint-mediated arrest.

## Discussion

Polyomaviruses rely on host DNA replication machinery to amplify their genomes, yet how viral replication interfaces with host cell-cycle progression at the single-cell level remains incompletely understood. Here we combined live-cell imaging, pharmacologic cell-cycle perturbation, flow cytometry, and single-cell transcriptomics to define the temporal relationship between Merkel cell polyomavirus (MCV) replication and host-cell-cycle dynamics. Across these orthogonal approaches we find that late viral gene expression, marked by VP1 accumulation, is not simply associated with proliferation but is instead gated to a window around late-S and G2 phases and followed by prolonged G2-like arrest, nuclear remodeling, and cell death. Live cell trajectories further show that VP1 expression typically begins after cells have entered S phase or G2, whereas cells that divide before detectable replication can transmit silent viral genomes to daughter cells that frequently initiate VP1 expression synchronously. Together, these findings support a model in which MCV genomes can persist in a pre-replicative silent state, then engage replication only when the host cell enters a permissive late-S/G2 state, coupling viral amplification to cell-cycle regulated non-mitotic cell fates.

The sister-cell analysis, though based on a modest cohort (12 lineages, 8 informative pairs) provides, to our knowledge, the first direct single-cell evidence that a polyomavirus genome can persist through mitosis in a transcriptionally silent state and subsequently reactivate synchronously in both daughter cells. The small sample size is an inherent consequence of this experimental design rather than a limitation of statistical power: identifying these events requires continuous single-cell tracking through both a pre-replicative mitosis and the subsequent days-long window of VP1 induction, the combination of two independently rare occurrences. We therefore interpret the high concordance between sisters (7 of 8 pairs within a 10-hour window) as a robust qualitative pattern rather than a precise estimate of penetrance, and the four asymmetric pairs as evidence that this inheritance is not absolute. Larger cohorts, achievable with extended imaging duration, and higher-throughput tracking, will be needed to define the quantitative penetrance of mitotic genome inheritance and to determine whether asymmetric outcomes reflect biological variation in genome copy number per cell versus detection-sensitivity limits of the reporter. This result expands the significance of MCV replication beyond cell-cycle timing because it suggests that viral genomes can remain replication competent across cell division and then reactivate coordinately in a cell lineage.

Live-cell imaging revealed that VP1 accumulation most often begins during late S phase or after entry into G2, while pharmacologic cell-cycle perturbation experiments demonstrated enrichment of VP1-positive cells when cultures were synchronized in G2. These findings were further supported by infection experiments and single-cell RNA sequencing in BJ-hTERT cells, where cells expressing both early and late viral transcripts were strongly enriched in S/G2 phases. Together, these complementary approaches indicate that late viral gene expression is strongly biased toward late S/G2 phases rather than occurring uniformly throughout the cell cycle. A single-cell analysis of BKPyV infection found that robust TAg early gene expression requires an initial host S phase^38^ ; given that VP1 protein accumulation necessarily follows a cascade of early gene expression, transcription, and translation, the late S/G2 window we observe for VP1 detection may reflect the same cell-cycle trigger measured further downstream.

Previous studies have observed that cells expressing VP1 fail to exit G2 phase in both BK polyomavirus (BKPyV) and SV40 polyomaviruses^39,40^. However, these studies did not directly observe in living single cells the state transition from pre-replicative to late gene positive replication as observed here. In our study, live cell imaging shows that MCV VP1+ cells have prolonged G2 phase, consistent with related polyomaviruses, and that VP1 accumulation occurs primarily during S and G2/M phases. The agreement between live-cell imaging, synchronization experiments, and BJ-hTERT infection data argues that cell-cycle coordination of VP1 expression is a core feature of the MCV life cycle. One possible explanation for this sustained G2 state is activation of replication-stress or DNA-damage checkpoint pathways during viral genome amplification. Polyomavirus large T antigens drive host replication machinery to initiate viral DNA synthesis. In this context, such replicative stress and host response pathways could be permissive and restrictive. Permissive by maintaining a replication-competent host environment, yet restrictive by preventing mitosis and promoting cell death in strongly replicating cells. Although the present study does not test ATR/Chk1-dependent checkpoint signaling, prolonged or arrested G2, enrichment of cell-cycle and DNA replication programs in ER+LR cells, and gradual loss of VP1+ cells is consistent with replication stress and checkpoint-engaged host responses.

Our orthogonal analyses in BJ-hTERT confirmed and extended our observations of cell-cycle timing and host-state remodeling. In both live cell imaging and single-cell RNA sequencing, VP1+ or ER+LR cells are enriched in S/G2 phases, supporting the conclusion that productive replication occurs preferentially after G1. At the transcriptional level, infected cultures mounted antiviral and inflammatory responses that were shared across all cells, together with a progressive downregulation of the cell shape and morphology genes as infection advanced. The decrease in cell-shape regulatory programs is consistent with the enlarged nuclei observed in VP1+ U2OS and BJ-hTERT cells. Throughout, replication, cell-cycle, and translational programs remain enriched in ER+LR cells. Together, these findings establish that MCV replication is coupled to a distinct host state characterized by late-S/G2 restriction, nuclear remodeling, and progressive loss of viability.

### Limitations of the study

A key limitation of this study was the duration over which live-cell dynamics could be continuously imaged before cell density and crowding altered cellular behavior. Reliable imaging was possible for approximately five days, which limited the ability to capture complete VP1 expression trajectories and terminal cell fates for many cells. As a result, some VP1-positive cells remained in extended S/G2 states without an observed outcome within the imaging window. Cell motility during long-term imaging also limited continuous single-cell tracking. This effect was minor in U2OS cells but more pronounced in BJ-hTERT cultures, where increased mobility and cell overlap restricted recovery of full time-course trajectories. Another limitation is that the live-cell imaging experiments monitored only late viral gene expression through VP1 reporters. The absence of fluorescent reporters for early gene expression limited direct comparison between imaging experiments and transcriptional states defined by single-cell RNA sequencing. Finally, single-cell RNA sequencing captures only viable cells present at the time of sampling and may therefore underrepresent transcriptional states associated with late-stage infection or cell death.

## Resource Availability

Additional information, as well as requests for resources and reagents, should be directed to Yuan Chang, Patrick S. Moore and Robin E. C. Lee.

## Materials availability

No unique reagents were generated in this study.

## Data and code Availability

All original code including the Live Cell Imaging Analysis (LCIA) toolbox has been deposited at Zenodo and is available at: https://doi.org/10.5281/zenodo.21349475

## Acknowledgments

We thank all the members of the Lee Lab and the Chang-Moore lab for the helpful discussions. This work was supported by the U.S. National Institutes of Health through grants R01 AI177607 (to Y.C). Additional support was provided by the Pittsburgh Foundation (to P.S.M.), the UPMC Foundation (to Y.C.), R35-GM119462 (to R.E.C.L.).

## Author contributions

S.W.S., H.Z., G.J.K., and D.L.S. prepared the cells and performed the live cell imaging experiments. S.W.S., H.Z., and G.J.K. performed the computational analysis of the live cell imaging experiment. B.A., Y.H. and M.M.H. performed and analyzed the single-cell sequencing experiments. S.W.S., H.Z., M.M.H., B.A., P.M., Y.C., and R.E.C.L. wrote the manuscript.

## Methods

### Cell culture

U2OS Geminin cells were generated by transfecting U2OS cells (ATCC HTB-96) with pCSII-EF-miRFP670v1-hGem(1/110) (Addgene plasmid #80006) and selecting with 500 µg/mL Zeocin. 293 TRE-sTco cells were generated by transducing 293 cells (ATCC CRL-1573) with pLenti TRE MCV sT and selecting with 2 µg/mL puromycin. BJ, BJ-hTERT, BJ-hTERT Geminin, 293, and 293 TRE-sTco cells were maintained in Dulbecco’s Modified Eagle Medium (DMEM; Corning, Manassas, VA, USA) supplemented with 10% fetal bovine serum (FBS). U2OS cells were maintained in McCoy’s 5A medium (Corning, Manassas, VA, USA) supplemented with 10% FBS.

BJ-hTERT cells were generated by retroviral transduction of Babe hTert-puro (pBabe-hTert-puro; a gift from Roderick J. O’Sullivan, Hillman Cancer Center, University of Pittsburgh, Pittsburgh, PA, USA) into primary BJ foreskin fibroblasts (ATCC CRL-2522), followed by selection with 2 µg/mL puromycin. BJ-hTERT Geminin cells were established by lentiviral transduction of CSII-EF-miRFP670v1-hGem(1/110) (Addgene plasmid #80006), followed by fluorescence-activated cell sorting.

### MCVmc transfection and infection

For MCVmc transfection, U2OS Geminin cells were plated in 6-well plates and transfected with 1 µg MCVmc-mScarlet using FuGENE (cat. no. E2311, Promega, Madison, WI, USA) according to the manufacturer’s instructions.

For production of fluorescent pseudo-MCV, 293 TRE-sTco cells were co-transfected in T75 flasks with 18 µg MCV genome DNA, 15 µg pWM plasmid (expressing VP1), and 5 µg ph2m plasmid (expressing VP2) and incubated for three days. For native MCv.mc Wt virion production, 293 TRE-sTco cells were co-transfected in 10 cm dish were transfected with 10 µg MCVmc.wt DNA and expanded into T175 flask in the presence of 5000 ng/ml Doxycycline for 10 days. Cells were harvested, virions were purified, and viral titers were quantified by RT-qPCR.

For MCV infection, BJ-hTERT Wt or Geminin stable cells were plated in 6-well plates and infected with purified MCVmc reporter or Wt virus (MOI = 10⁴) or mock in the presence of epidermal growth factor (EGF, 10 µg/mL), basic fibroblast growth factor (bFGF, 20 ng/mL), CHIR9901 (3 µM), and collagenase IV (1 mg/mL).

### Live cell Imaging

On the day of the experiment, growth medium was aspirated from each well of the 96-well plate and replaced with imaging medium containing trace amounts of Hoechst (50 ng/mL). Imaging was performed in FluoroBrite DMEM medium (Gibco, A18967-01) supplemented with 10% FBS (Corning), penicillin (100 U/mL), streptomycin (100 U/mL), and 0.2 mM L-glutamine. Images were taken every 15 minutes over multiple days and was performed on a DeltaVision Elite microscope (GE Healthcare) fitted with a pco.edge sCMOS camera, an Insight solid-state illumination module, and a 20X objective, within an environmentally controlled chamber maintained at 37 °C and 5% CO₂.

### Image analysis

Nuclear segmentation of Hoechst-stained images was performed using the open-source deep learning algorithm CellPose v3.1.0^41^. An initial cell diameter was estimated automatically from representative images in the graphical interface. The built-in nuclei model was then fine-tuned on our own sample images, where segmentation errors were manually corrected, and the model was retrained through the GUI to improve accuracy. Separate models were trained for U2OS and BJ-hTERT cells to ensure optimal segmentation performance for each cell type. A custom Python pipeline was used to segment and save each frame automatically on a workstation equipped with an Intel® Xeon® E5-2630 v4 CPU (2.20 GHz) and an NVIDIA GeForce GTX 1080 GPU (8 GB). GPU acceleration in CellPose was enabled to increase segmentation efficiency.

Single-cell trajectories were generated using bTrack v0.6.5^42^, which automatically detected and linked cells across frames from the segmentation mask generated by CellPose. The default settings were used, and the max search radius was set to 50 pixels. Since VP1 expression events are relatively rare, precise single-cell trajectories were essential. To address this, a custom Napari-based Python tool was created that enabled manual correction of tracking errors and annotation of mitosis and cell death events. Therefore, every cell trajectory was manually confirmed and corrected to ensure accuracy. The code is available at https://github.com/recleelab/LCIA.

Three fluorescent channels were acquired during live-cell imaging: Hoechst, VP1– mScarlet, and miRFP670–hGem(1–110). Each frame was background-corrected by subtracting the pixel intensity mode. VP1–mScarlet images were additionally flat-field corrected because signal intensities were near the noise floor. For each frame, nuclear masks were used to compute per-cell statistics, including mean and integrated nuclear intensity. Single-cell trajectories were generated from manually verified tracks, with each mitotic event initiating a new trajectory.

Cells were classified as VP1(+) or Geminin (+) based on sustained reporter expression. A cell was defined as VP1(+) if its integrated nuclear VP1–mScarlet intensity exceeded the 99th percentile of the VP1(–) population for at least one consecutive hour during imaging. This threshold minimized false positives resulting from random fluctuations or transient increases in fluorescence due to imaging conditions. Geminin(+) cells were identified using the same criterion, substituting mean nuclear intensity of the Geminin reporter for VP1–mScarlet.

### Cell cycle arrest and flow cytometry

U2OS cells were transfected as described above. Three days post-transfection, cells were treated with 0.5 mM mimosine, 10 µM aphidicolin, 9 µM RO3306, or vehicle control for 24 h. Cells were then released from treatment and pulsed with 100 µM BrdU for 30 min before harvesting.

For cell cycle phase determination by flow cytometry, cells were stained intracellularly with mouse anti-BrdU Alexa Fluor 488 (1:10 dilution, CM Log# 301) and FxCycle Violet Kit (cat. no. F10347, Thermo Fisher Scientific, Waltham, MA, USA). Flow cytometry was performed on an LSRFortessa (BD Biosciences).

### Single-cell RNA-seq data processing

Single-cell RNA-sequencing (scRNA-seq) data were generated using the 10x Genomics Chromium Single Cell 3′ Gene Expression platform^43^ with multiplexed samples collected at 3, 5, and 7 days post-infection (dpi) from Merkel cell polyomavirus (MCV)-infected and mock-treated conditions.

### Read Alignment and Quantification

The raw sequencing data were processed using the Cell Ranger software suite (10x Genomics, version 7.0) with the standard pipeline. For read alignment and gene quantification, a custom reference genome was constructed by concatenating the human genome (GRCh38/hg38) and the MCV viral genome (GenBank accession JF813003.1)^28^. Gene annotations for human transcripts were obtained from the Ensembl GRCh38 release, and two viral genes corresponding to the early and late MCV transcriptional units were appended to the reference annotation file. Cell Ranger was used to perform demultiplexing, alignment, barcode processing, and unique molecular identifier (UMI) counting following the manufacturer’s standard workflow. The resulting gene-by-cell count matrices were generated for each sample and used for downstream quality control and analysis.

### Quality Control

Cells with low-quality or potentially artifactual profiles were removed prior to downstream analysis (Supplementary Fig. 6D,E). Specifically, cells were retained if they had more than 200 but fewer than 7,500 detected genes (nFeature_RNA), and more than 500 but fewer than 60,000 total UMI counts (nCount_RNA). Cells with high mitochondrial gene expression (>15%) were excluded to remove stressed or dying cells. These filtering criteria ensured removal of low-quality cells, potential doublets, and extreme outliers while retaining high-quality single-cell transcriptomes for downstream analyses.

### Identification of MCV-Infected Cells

To identify MCV-infected cells, we quantified expression of the viral early and late genes in single cells. Cells were considered infected if they had more than 1 UMI for the early gene and more than 2 UMIs for the late gene. These thresholds were chosen to ensure statistically confident detection of infection, as a small number of viral reads were observed in mock-treated control cells. To further justify these cutoffs, we computed the 99th percentile of viral gene expression in mock samples, and only cells exceeding this level were classified as infected. This approach minimizes false-positive infection calls while capturing confidently infected cells for downstream analyses.

### Cell cycle assignment to cells

Cell cycle phase assignment was performed using Seurat’s^44^ CellCycleScoring function. Prior to scoring, gene expression counts were normalized and variance-stabilized using the SCTransform function with default parameters. Canonical marker genes for S and G2/M phases were used to assign each cell to the corresponding phase. Cells that did not meet the criteria for S or G2/M phases were classified as G1 by default.

### Differential expression analysis

Differential gene expression analysis between two groups (e.g., infected vs. mock or bystander vs. mock) was performed using Seurat’s FindMarkers function. The Wilcoxon rank-sum test (test.use = "wilcox") was applied to assess statistical significance. To control for biases due to unequal cell numbers, cells were randomly downsampled so that both groups contained the same number of cells as the smallest group, ensuring that detected differences in gene expression reflected biological variation rather than group size.

### Functional enrichment analysis

Functional enrichment analysis was performed using the gseGO function from the clusterProfiler R package^45^. Differentially expressed genes were ranked by log-fold change and used as input (geneList). Gene Ontology (GO) enrichment analysis was conducted for the Biological Process (BP) category using the org.Hs.eg.db annotation database. Gene sets with sizes between 20 and 500 genes were considered, and statistical significance was assessed with a *p* value cutoff of 0.05. P-values are computed based on the gene set enrichment analysis (GSEA) statistical framework, which is a Kolmogorov–Smirnov (KS)-like test. P-values were adjusted for multiple testing using the Benjamini–Hochberg method (pAdjustMethod = "BH").

### Transcription factor analysis

We inferred transcription factor activity from normalized single-cell RNA expression profiles using the human DoRothEA TF-target regulon resource together with the VIPER method as implemented in decoupleR. TF activity scores were computed across all retained cells using run_viper function on the normalized RNA assay, restricting analysis to regulons with at least 10 target genes to improve scoring stability. For statistical comparison of inferred TF activities between groups, the VIPER-derived TF activity matrix was added back into Seurat as a dedicated TF_activity assay, and group differences were tested using Seurat’s Wilcoxon rank-sum framework (FindMarkers) on TF activity values. For imbalanced comparisons, including ER+LR versus mock, we used an adaptive repeated-downsampling procedure: the larger group was randomly subsampled to match the smaller group for 20 iterations, Wilcoxon testing was repeated in each iteration, and adjusted p-values were aggregated using Fisher’s method. A TF was considered robustly differential only if it satisfied all of the following: combined p-value < 0.05, significance in at least 80% of iterations, consistent direction of effect across iterations, and absolute mean log2 fold-change in inferred activity >= 0.3.

