## Supplemental for "Cell Cycle Regulation of Merkel Cell Polyomavirus Replication and Genome Inheritance in Single Cells"

**Other footnotes**

[*]Corresponding Authors

**Supplementary figures**

**Supplementary Figure 1: Threshold determination of nuclear Geminin and VP1 postive cells**

A. Histogram of the mean flourescent intensity (MFI) of the nuclear geminin of the mock (blue) and transfected (orange) cells. B. Histogram of the integrated flourescent intensity (IFI) of the mock (blue) and transfected (orange) cells. For both A and B, the 99$^{th}$ percentile of the mock cells served as the threshold for expression. C. Bar plot showing the percent of cells with observed cell death classified as being Mock, VP1-, and VP1+ cells. Fisher exact test used to determine significance.

Figure S1

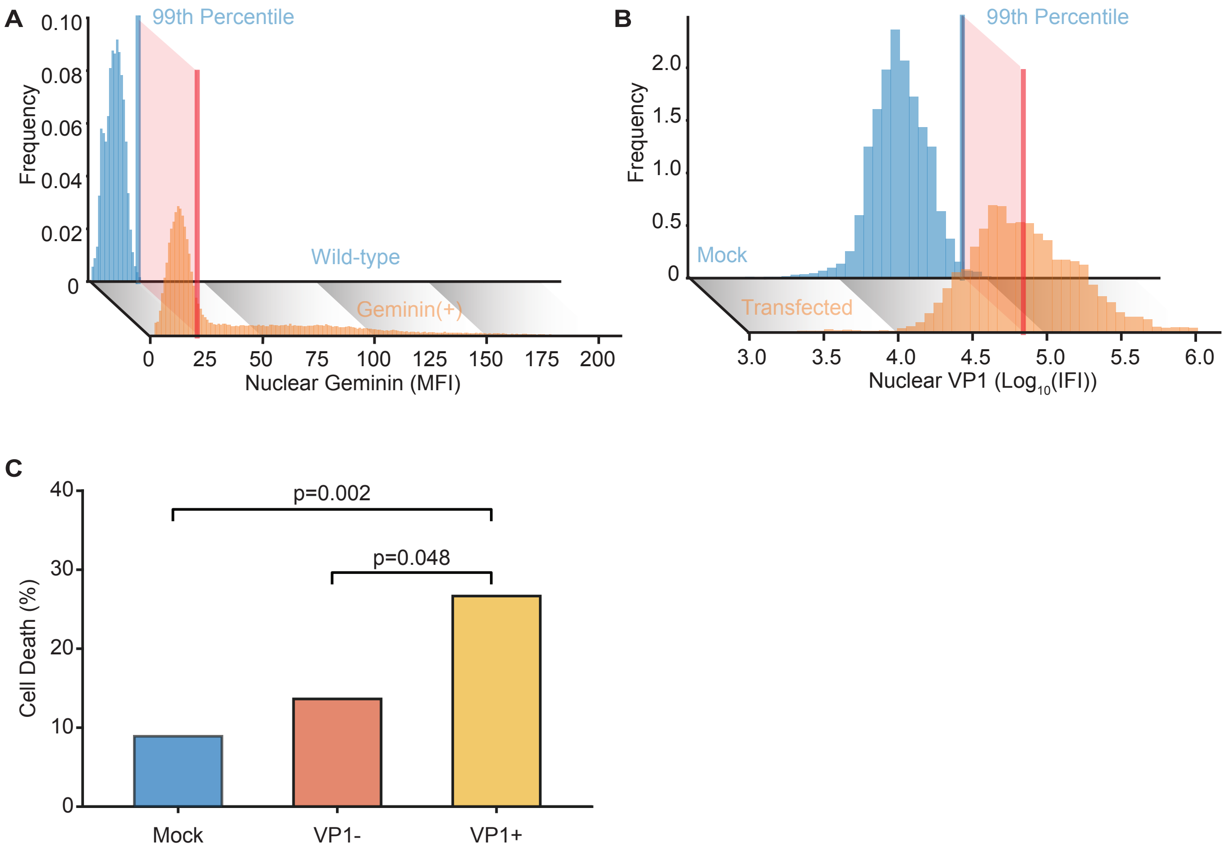

A
0.10
0.08
0.06
0.04
0.02
0
Frequency
99th Percentile
Wild-type
Geminin(+)
0
25
50
75
100
125
150
175
200
Nuclear Geminin (MFI)
B
2.0
1.5
1.0
0.5
0
Frequency
99th Percentile
Mock
Transfected
3.0
3.5
4.0
4.5
5.0
5.5
6.0
Nuclear VP1 (Log10(IFI))
C
40
30
20
10
0
Cell Death (%)
p=0.002
p=0.048
Mock
VP1-
VP1+

**Supplementary Figure 2: Heatmap of the mock and VP1- cells defining the start of each phase**

A. Table indicating what each class represents as the obserbable with either observed mitotic birth or fate. Red X define cell death and green circles represent mitosis. B. Kymographs of mock and VP1- U2OS cell trajectories, separated into the four classes. Within each class, trajectories are aligned by the start of S phase. Each trajectory displays mean fluorescence intensity (MFI) of nuclear Geminin, integrated fluorescence intensity (IFI) of nuclear Hoechst, and nuclear VP1 IFI over time. Rightward arrows mark the start of S phase, and leftward arrows mark the start of G2, shown as white in the Geminin MFI or Hoechst IFI plots, and as red or blue, respectively, in the VP1 IF. (C) Boxplot showing the duration of geminin positivity (hours) across the trajectory classes defined in panel A. Each class includes Mock, VP1(–), and VP1(+) groups, with sample sizes of 16, 10, and 37 (Class 1); 17, 26, and 16 (Class 2); 42, 29, and 14 (Class 3); and 26, 23, and 8 (Class 4), respectively. Horizontal bars indicate statistically significant pairwise differences between groups as determined by two-sample t-tests.

Figure S2

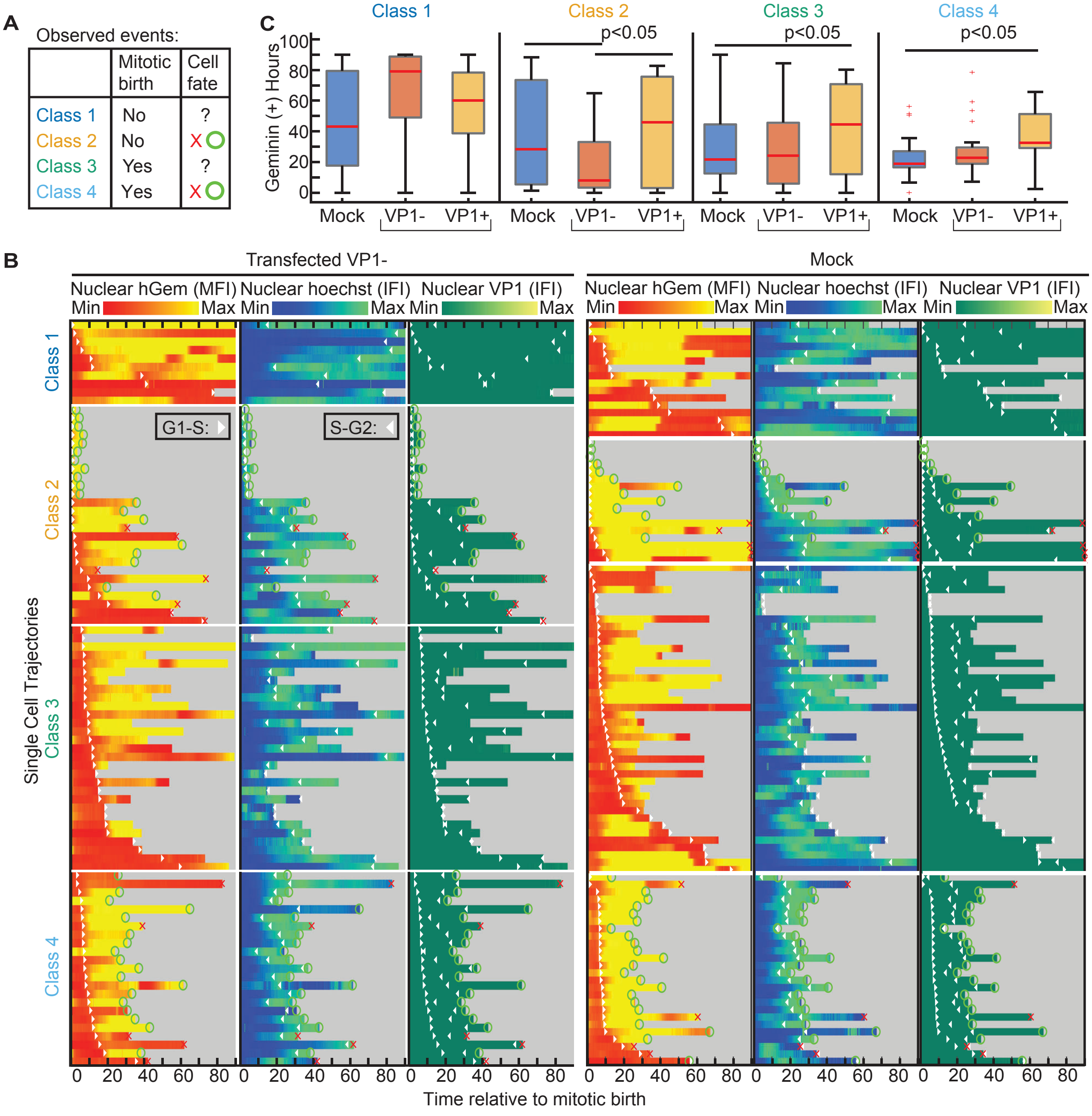

A
Observed events:
Mitotic birth
Cell fate
Class 1 No ?
Class 2 No X O
Class 3 Yes ?
Class 4 Yes X O
C
Class 1
Class 2
Class 3
Class 4
p<0.05
p<0.05
p<0.05
Geminin (+) Hours
100
80
60
40
20
0
Mock
VP1-
VP1+
B
Transfected VP1-
Mock
Nuclear hGem (MFI)
Nuclear hoechst (IFI)
Nuclear VP1 (IFI)
Min
Max
G1-S:
S-G2:
Single Cell Trajectories
Class 1
Class 2
Class 3
Class 4
0 20 40 60 80
Time relative to mitotic birth

**Supplementary Figure 3: Symmetric and asymmetric VP1 expression in daughter cells**

Example images of daughter cells following mitosis showing (A) symmetric fate, where both daughters express VP1, and (B) asymmetric fate, where only one daughter expresses VP1. White arrows mark VP1-negative cells, and red arrows mark VP1-positive cells.

Figure S3

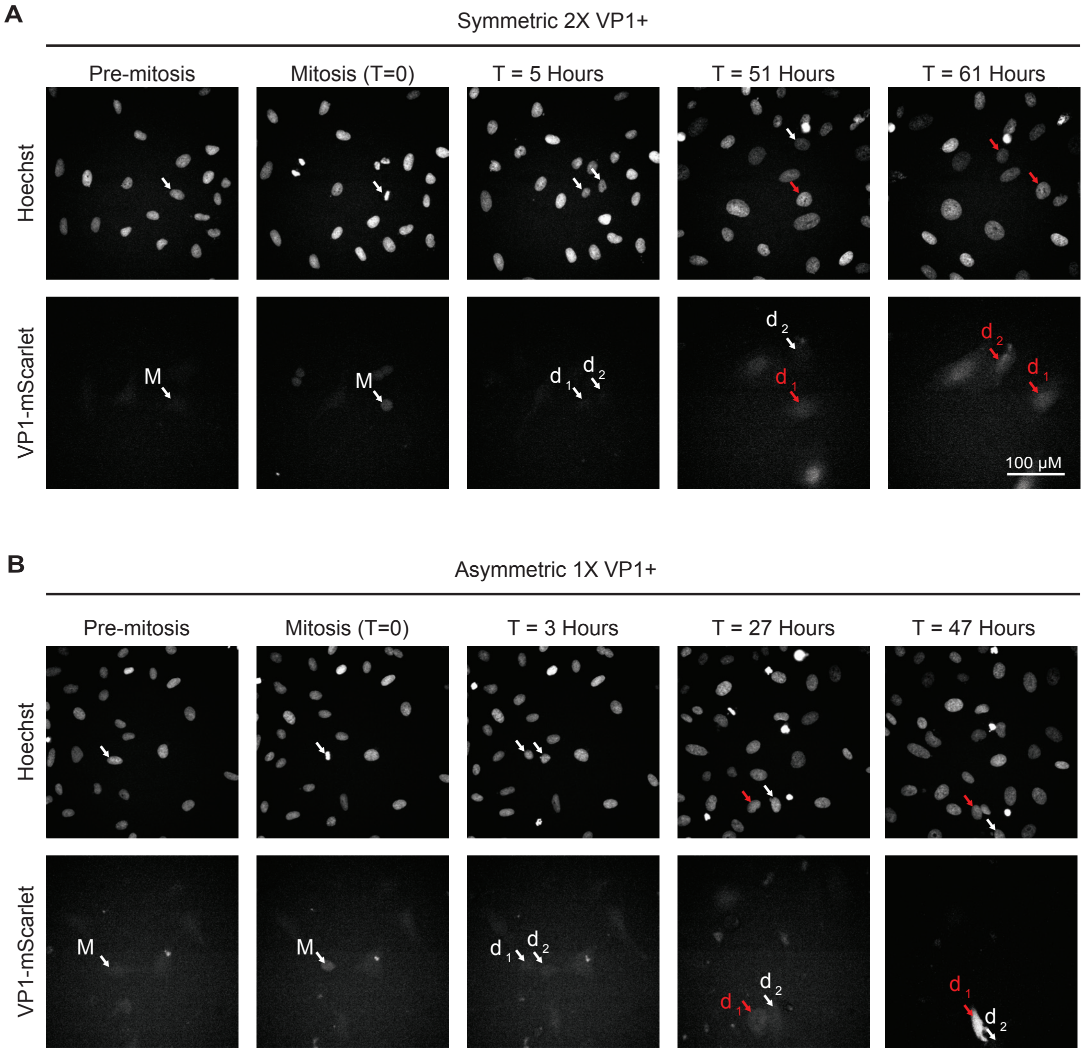

A
Symmetric 2X VP1+
Pre-mitosis
Mitosis (T=0)
T = 5 Hours
T = 51 Hours
T = 61 Hours
Hoechst
VP1-mScarlet
M
d 1
d 2
100 μM
B
Asymmetric 1X VP1+
Pre-mitosis
Mitosis (T=0)
T = 3 Hours
T = 27 Hours
T = 47 Hours
Hoechst
VP1-mScarlet

**Supplementary Figure 4: Cell-cycle progression following release from drug-induced arrest.**

A. Flow cytometry analysis of BrdU incorporation (y-axis) versus DNA content (x-axis) in U2OS cells at 0, 6, 12, and 24 hours after release from cell cycle inhibitors, which included unarrested, mimosine, aphidolin, and RO-3306. Quadrants indicate G1 (Q4, low BrdU), early/late S (Q1–Q2, high BrdU), and G2/M (Q3, low BrdU, 4N DNA). B. Heatmap displaying the conditional probabilty of a cell expressing VP1 given the cell cycle and time since cell arrestor release. The white represents condition with limited cells to make an accurate estimate. C. Bar plot showing the percentage of VP1-positive cells at 0, 6, 12, and 24 hours after release from each cell cycle inhibitor.

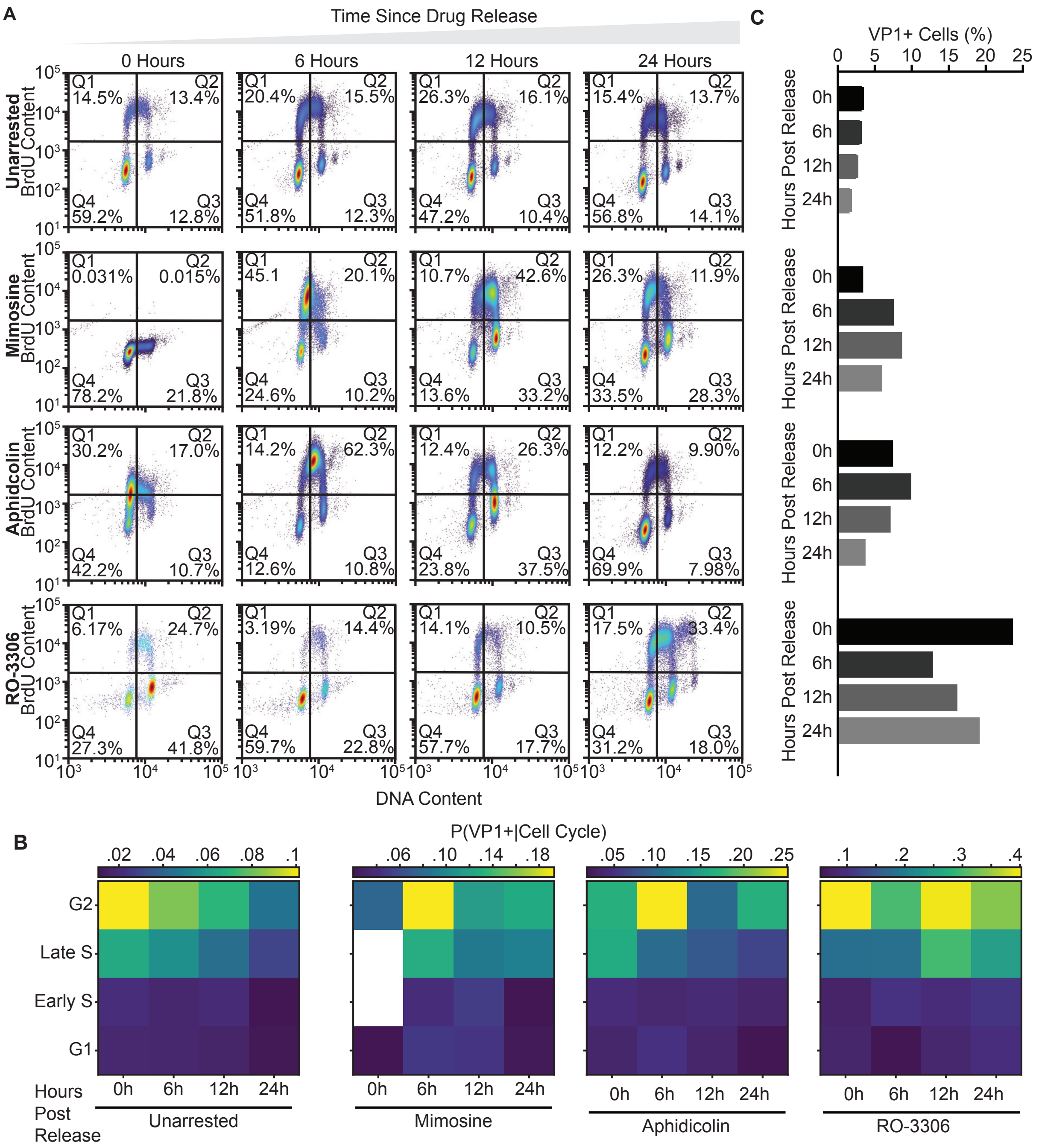
A
Time Since Drug Release
0 Hours
6 Hours
12 Hours
24 Hours
Unarrested
Mimosine
Aphidcolin
RO-3306
BrdU Content
DNA Content
Q1 14.5% Q2 13.4% Q4 59.2% Q3 12.8%
Q1 20.4% Q2 15.5% Q4 51.8% Q3 12.3%
Q1 26.3% Q2 16.1% Q4 47.2% Q3 10.4%
Q1 15.4% Q2 13.7% Q4 56.8% Q3 14.1%
Q1 0.031% Q2 0.015% Q4 78.2% Q3 21.8%
Q1 45.1 Q2 20.1% Q4 24.6% Q3 10.2%
Q1 10.7% Q2 42.6% Q4 13.6% Q3 33.2%
Q1 26.3% Q2 11.9% Q4 33.5% Q3 28.3%
Q1 30.2% Q2 17.0% Q4 42.2% Q3 10.7%
Q1 14.2% Q2 62.3% Q4 12.6% Q3 10.8%
Q1 12.4% Q2 26.3% Q4 23.8% Q3 37.5%
Q1 12.2% Q2 9.90% Q4 69.9% Q3 7.98%
Q1 6.17% Q2 24.7% Q4 27.3% Q3 41.8%
Q1 3.19% Q2 14.4% Q4 59.7% Q3 22.8%
Q1 14.1% Q2 10.5% Q4 57.7% Q3 17.7%
Q1 17.5% Q2 33.4% Q4 31.2% Q3 18.0%
C
VP1+ Cells (%)
0 5 10 15 20 25
Hours Post Release
0h 6h 12h 24h
B
P(VP1+|Cell Cycle)
.02 .04 .06 .08 .1
.06 .10 .14 .18
.05 .10 .15 .20 .25
.1 .2 .3 .4
G2
Late S
Early S
G1
Hours Post Release
0h 6h 12h 24h
Unarrested
Mimosine
Aphidcolin
RO-3306

**Supplementary Figure 5: Nuclear morphology and cell-cycle distribution of BJhTert MCV-infected cells**

A. Boxplots showing BJhTert nuclear area in mock, VP1(–), and VP1(+) cells at 5 and 7 days post-infection in live cell imaging experiments. VP1(+) cells exhibited significantly larger nuclei compared to uninfected or VP1(–) cells at both time points. A one-sided T-test determined statistical significance. B. Stacked bar plots showing the proportion of cells in G1, S, and G2/M phases at 5 and 7 days post-infection based on single-cell RNA-seq data. Cells were grouped as mock or MCV-infected, with infected cells further classified as bystander (no viral transcripts), only early gene–expressing, late gene–expressing, or infected (expressing either early or late genes).

Figure S5

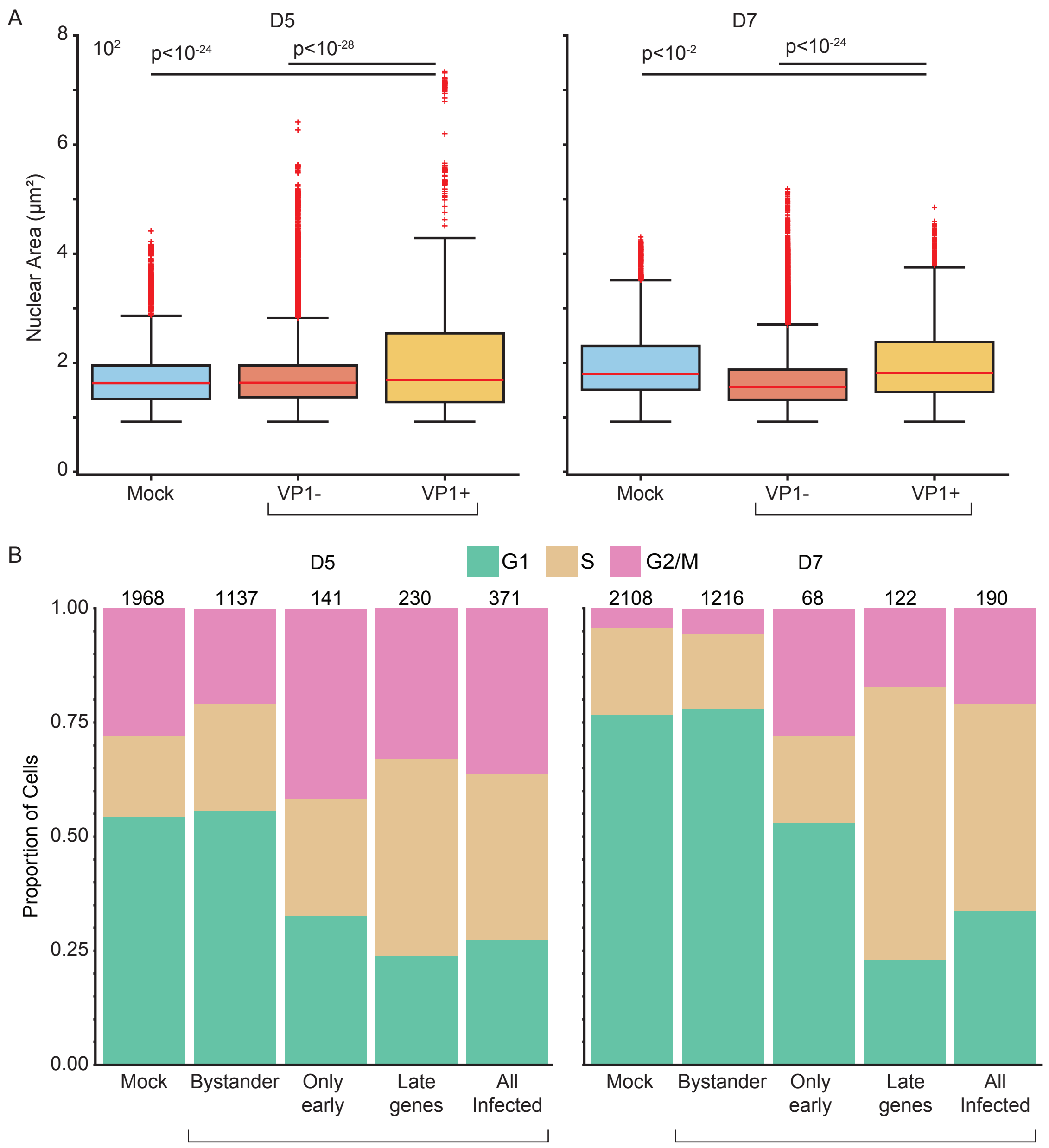
A
D5
D7
Nuclear Area (μm²)
10²
p<10⁻²⁴
p<10⁻²⁸
p<10⁻²
p<10⁻²⁴
8
6
4
2
0
Mock
VP1-
VP1+
B
D5
D7
G1
S
G2/M
1968
1137
141
230
371
2108
1216
68
122
190
Proportion of Cells
1.00
0.75
0.50
0.25
0.00
Mock
Bystander
Only early
Late genes
All Infected

**Supplementary Figure 6. Differential expression and quality control across single-cell viral infection states.**

A. Number of differentially expressed genes (DEGs) (padj < 0.05, |log2FC| > 1) per pairwise comparison at Day 5 (top) and Day 7 (bottom). Columns are defined by the transcriptional class of the test population (early-/late-gene positive) and the reference population (Vs.: Mock, Bystander, or early-gene-positive; grayscale shading). B. Volcano plot of differential inferred transcription factor (TF) activity in ER+LR cells at Day 7 versus Day 5. TF activity scores were inferred from target-gene expression using VIPER with DoRothEA regulons, and differential testing was performed on those scores. Each point represents one TF; positive x-axis values indicate higher inferred TF activity at Day 7, and negative values indicate higher inferred TF activity at Day 5. The y-axis shows -log10 adjusted P value. TFs meeting the significance and effect-size thresholds (adjusted P < 0.05, |activity difference| >= 0.3) are labeled. C. Similar to Figure 1B, volcano plots of differential inferred transcription factor (TF) activity in ER+LR cells, here comparing ER+LR cells with mock cells at Day 5 and Day 7. D. Per-cell QC metrics across the dataset: RNA counts (UMIs), detected genes, and % mitochondrial counts. Points, single cells; violins, distributions. E. QC metrics as in D, split by timepoint (Day 3/5/7) and infection status (checkmark, infected).

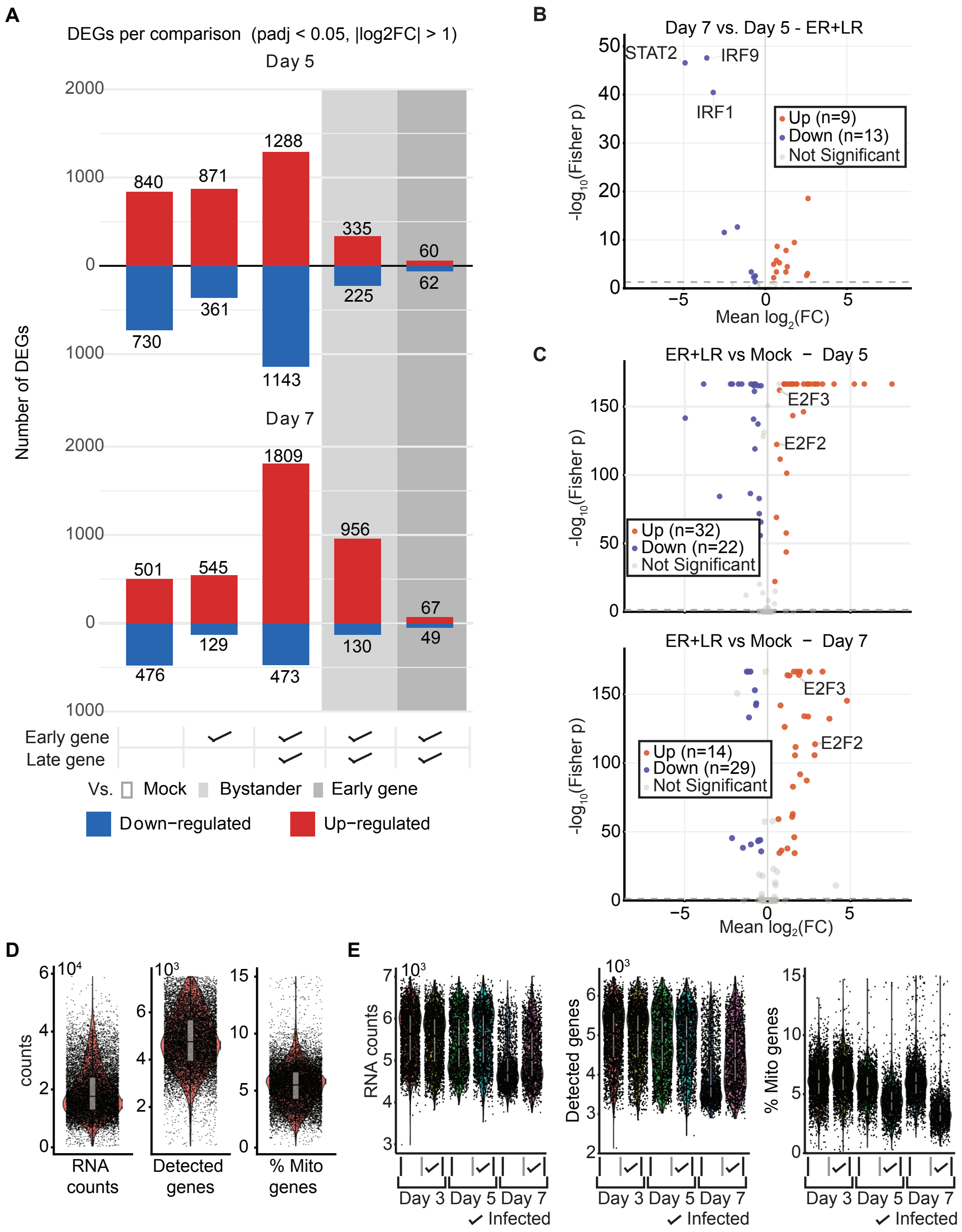
A
DEGs per comparison (padj < 0.05, |log2FC| > 1)
Day 5
Day 7
Number of DEGs
Early gene
Late gene
Vs. Mock Bystander Early gene
Down−regulated Up−regulated
B
Day 7 vs. Day 5 - ER+LR
STAT2
IRF9
IRF1
Up (n=9)
Down (n=13)
Not Significant
Mean log2(FC)
-log10(Fisher p)
C
ER+LR vs Mock − Day 5
E2F3
E2F2
Up (n=32)
Down (n=22)
Not Significant
ER+LR vs Mock − Day 7
Up (n=14)
Down (n=29)
Not Significant
D
RNA counts
Detected genes
% Mito genes
E
RNA counts
Detected genes
% Mito genes
Day 3 Day 5 Day 7
Infected

**Supplementary tables**

**Supplementary Table 1. Differential expression results for curated gene sets across infection-state comparisons.**

Grouped differential expression summary for genes assigned to nine curated biological programs across Day 5 and Day 7 comparisons. For each gene within each program, the table reports the comparison label, direction of change (up, down, or none), average log2 fold change (avg_log2FC), adjusted p value (p_val_adj), significance flag (sig_flag), and significance score (sig_score) for the following contrasts: bystander vs mock, ER vs mock, ER+LR vs mock, ER+LR vs ER, and ER+LR vs bystander.
